## Supplementary figures and images for "Polypyrimidine Tract Binding Proteins are essential for B cell development"

### Supplemental Figures

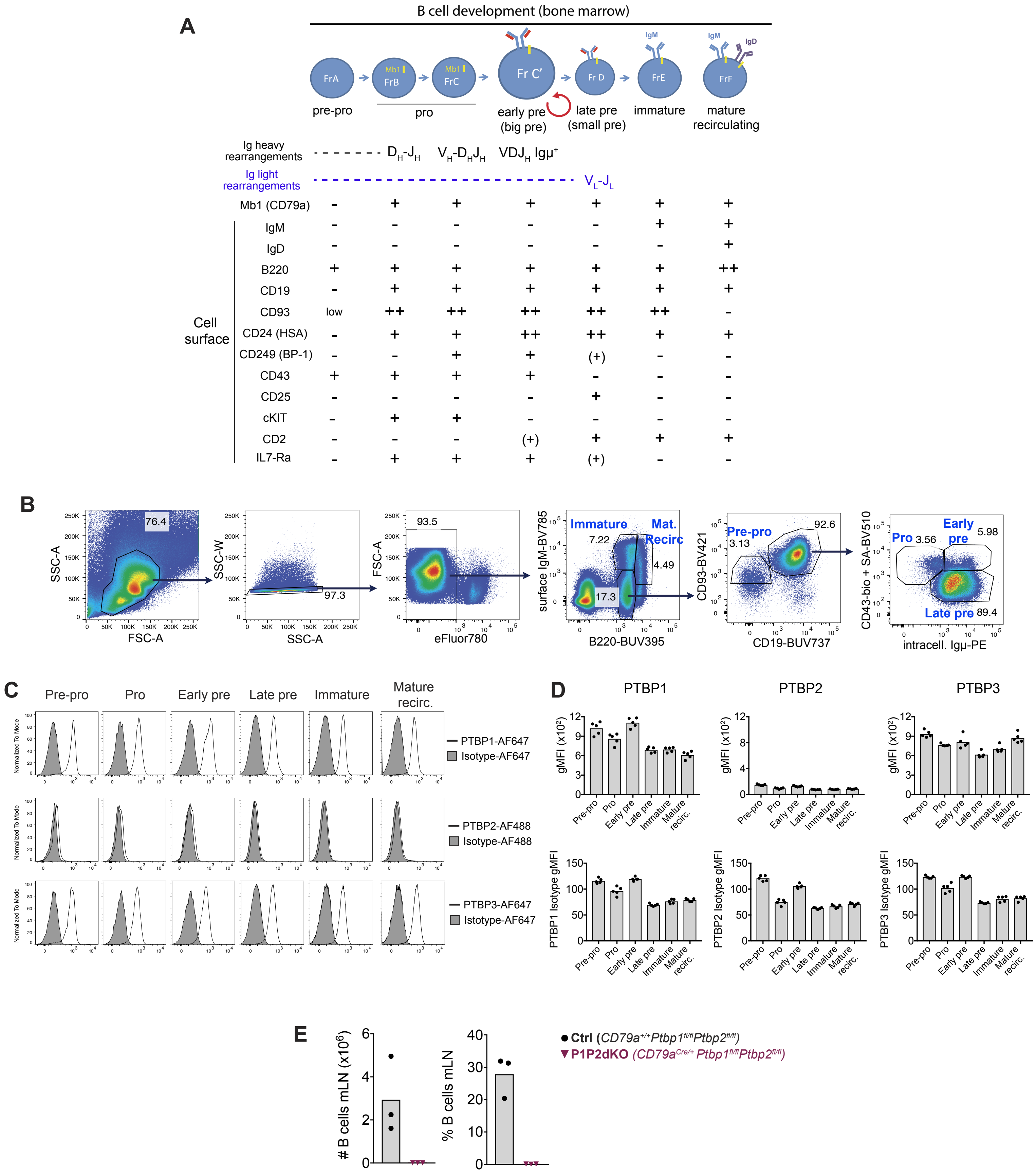

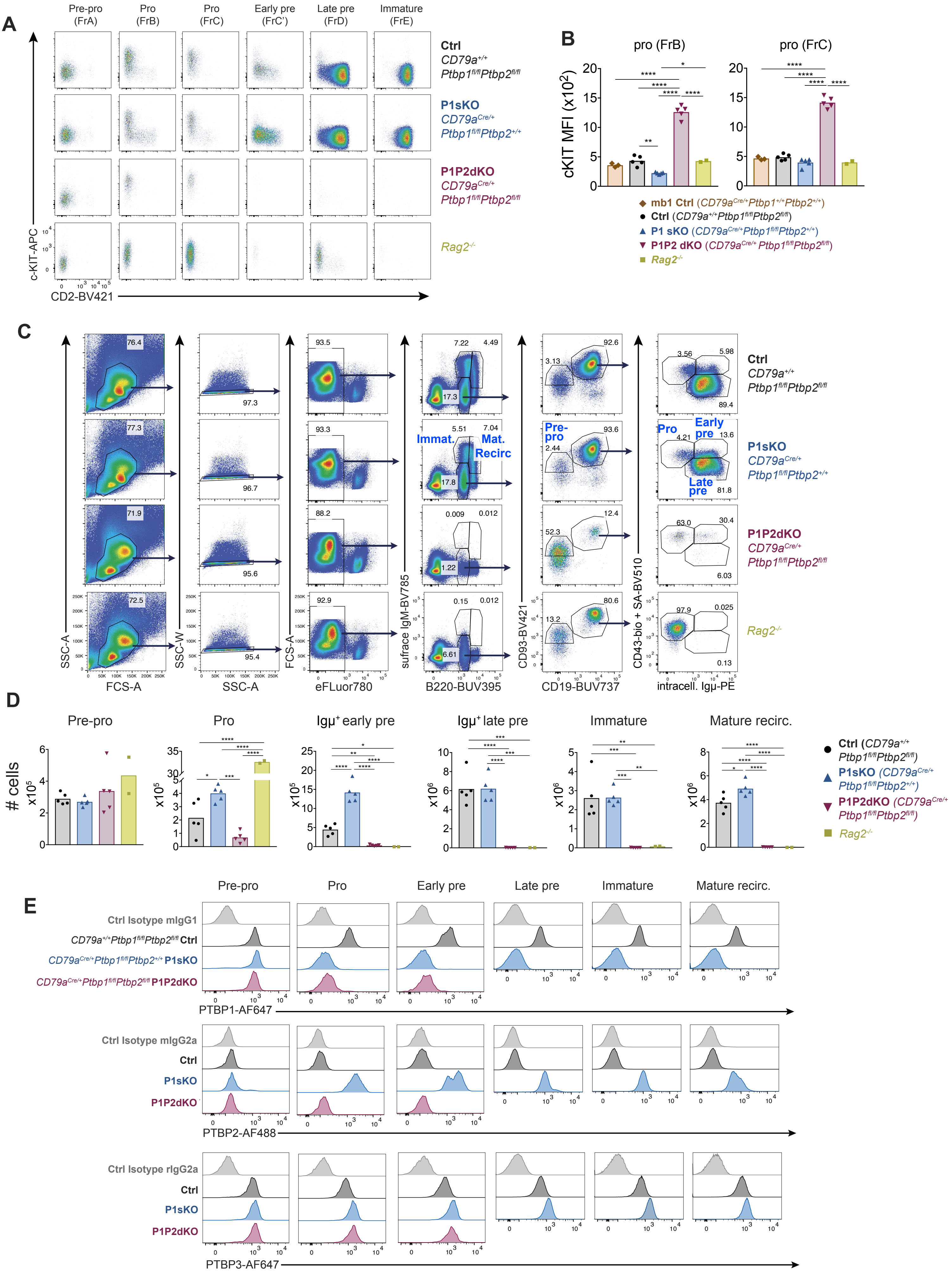

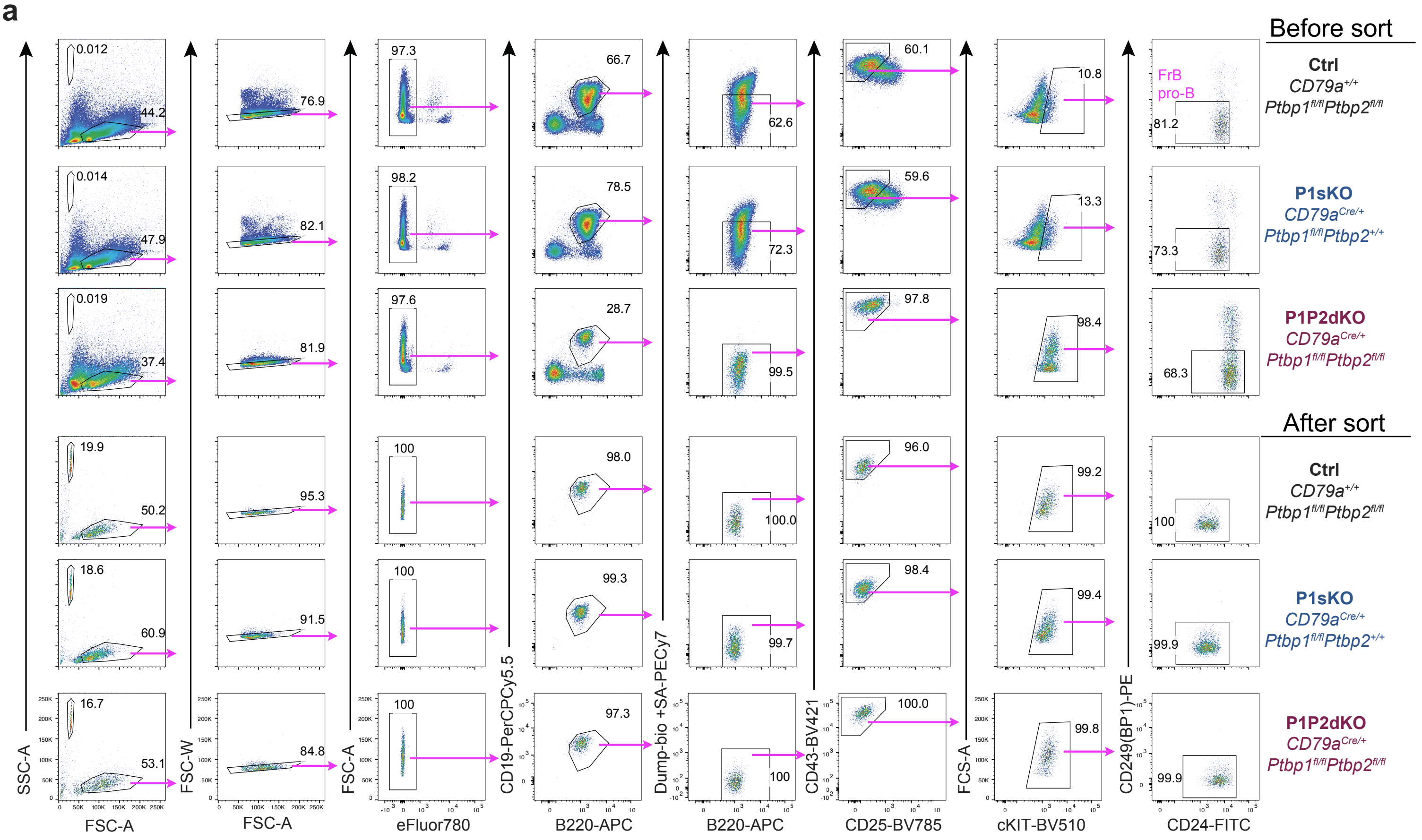

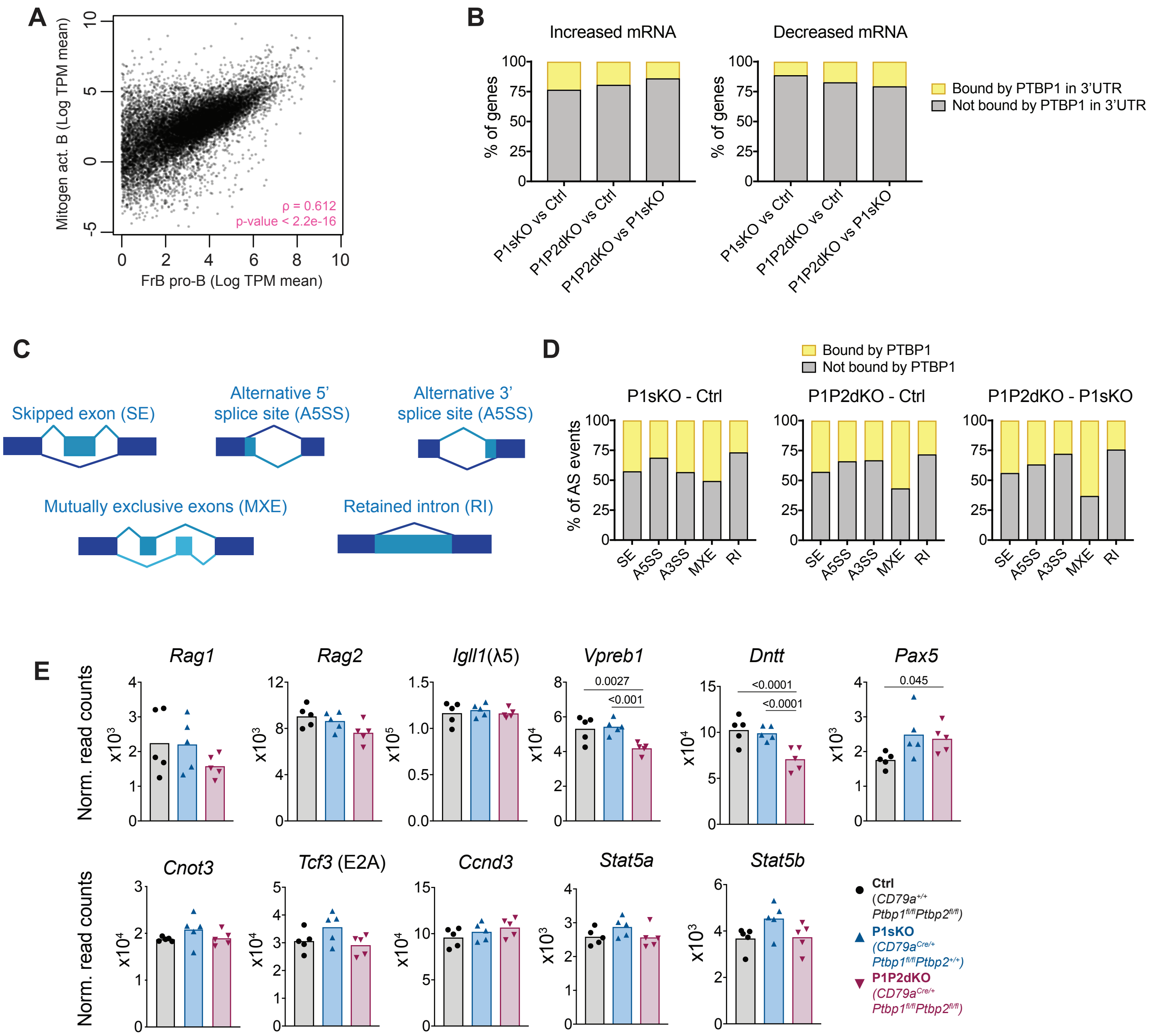

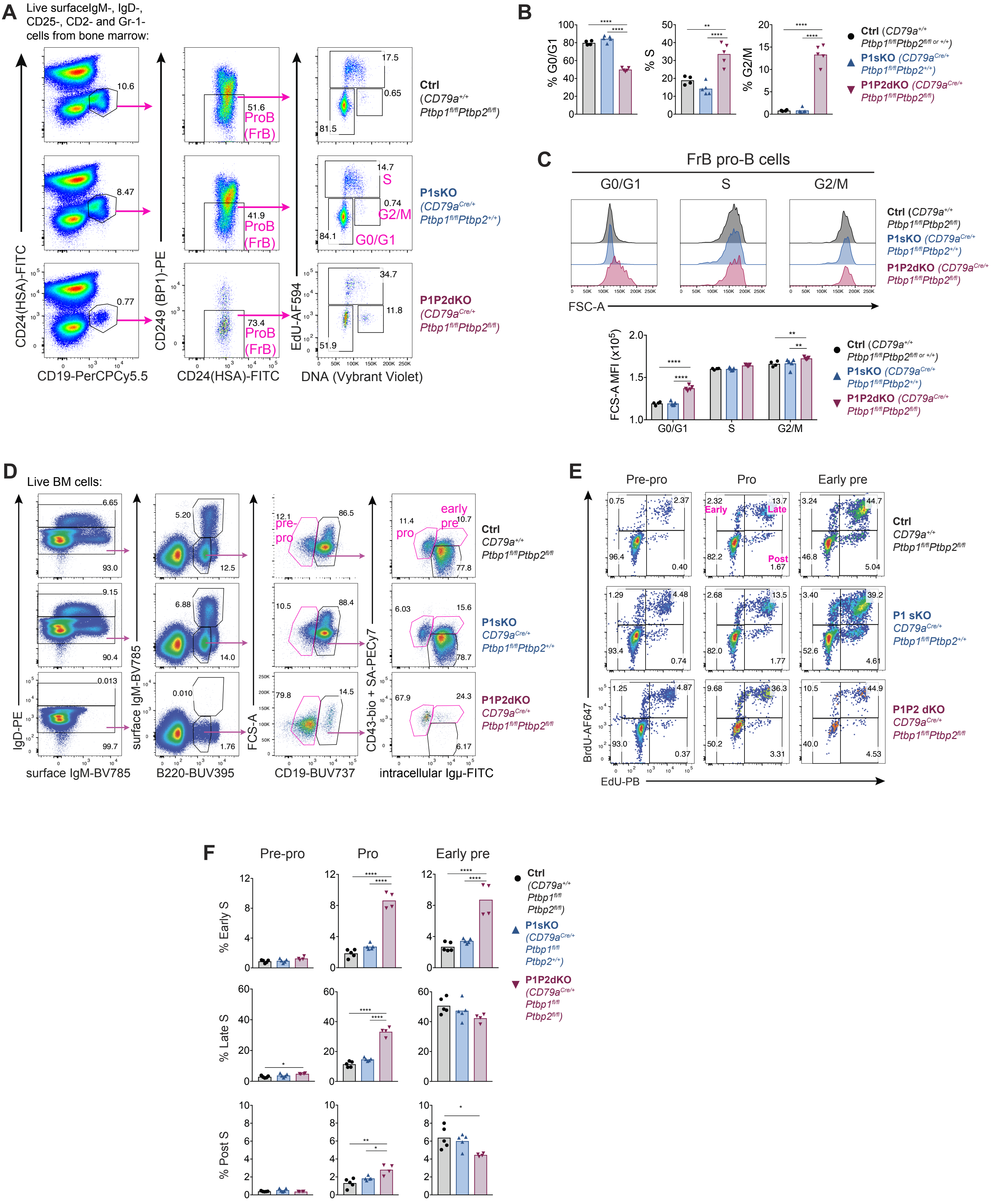

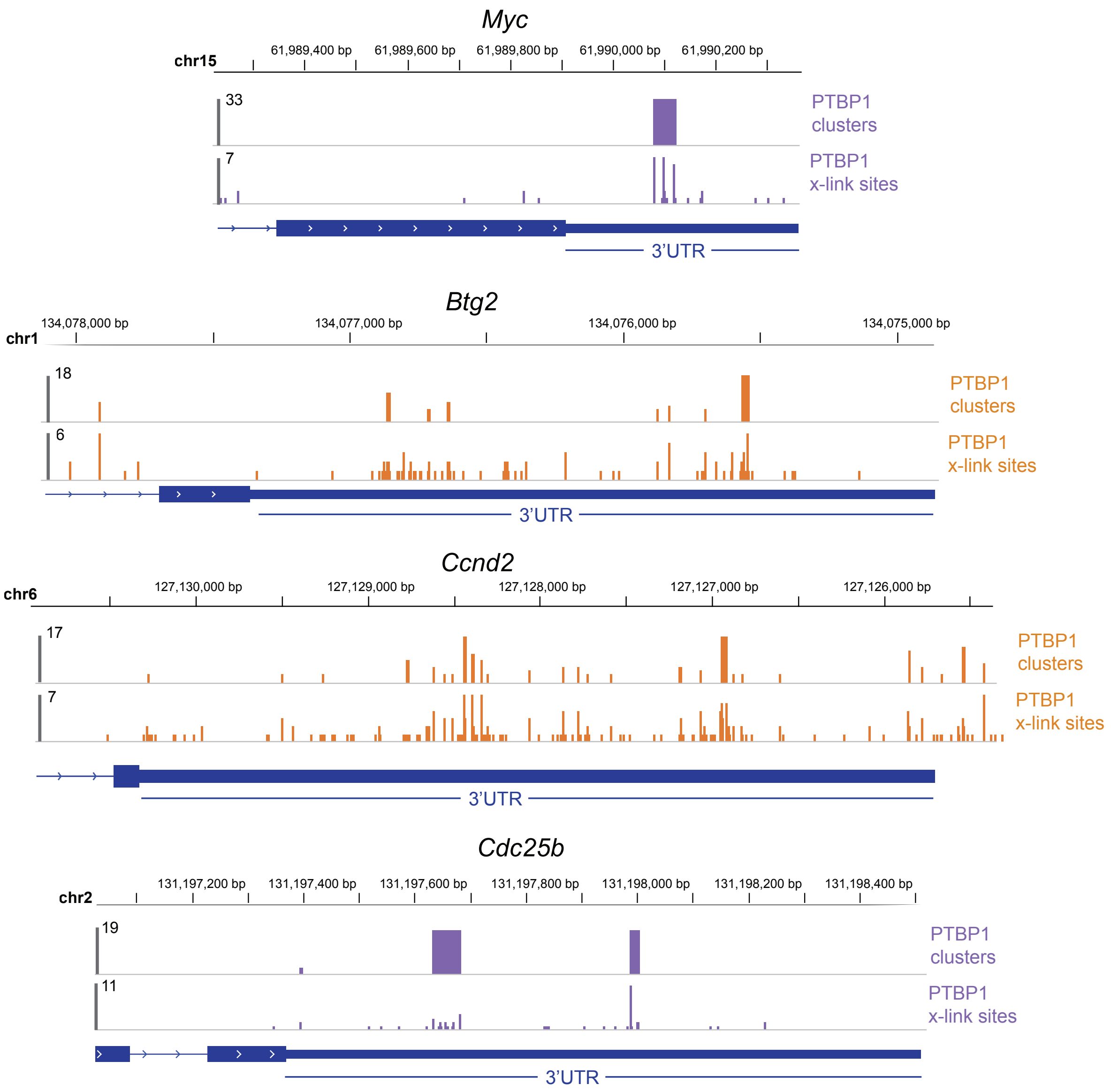
